# Hidden viral sequences in public sequencing data and warning for future emerging diseases

**DOI:** 10.1101/2021.05.17.444395

**Authors:** Junna Kawasaki, Shohei Kojima, Keizo Tomonaga, Masayuki Horie

## Abstract

RNA viruses cause numerous emerging diseases, mostly due to transmission from mammalian and avian reservoirs. Large-scale surveillance of RNA viral infections in these animals is a fundamental step for controlling viral infectious diseases. Metagenomic analysis is a powerful method for virus identification with low bias and has substantially contributed to the discovery of novel viruses. Deep sequencing data have been collected from diverse animals and accumulated in public databases, which can be valuable resources for identifying unknown viral sequences. Here, we screened for infections of 33 RNA viral families in publicly available mammalian and avian sequencing data and found approximately 900 hidden viral infections. We also discovered six nearly complete viral genomes in livestock, wild, and experimental animals: hepatovirus in a goat, hepeviruses in blind mole-rats and a galago, astrovirus in macaque monkeys, parechovirus in a cow, and pegivirus in tree shrews. Some of these viruses were phylogenetically close to human pathogenic viruses, suggesting the potential risk of causing disease in humans upon infection. Furthermore, the infections of five novel viruses were identified in several different individuals, indicating that their infections may have already spread in the natural host population. Our findings demonstrate the reusability of public sequencing data for surveying viral infections and identifying novel viral sequences, presenting a warning about a new threat of viral infectious disease to public health.

**Importance:** Monitoring the spread of viral infections and identifying novel viruses capable of infecting humans through animal reservoirs are necessary to control emerging viral diseases. Massive amounts of sequencing data collected from various animals are publicly available, and these data may contain sequences originating from a wide variety of viruses. Here, we analyzed more than 46,000 public sequencing data and identified approximately 900 hidden RNA viral infections in mammalian and avian samples. Some viruses discovered in this study were genetically similar to pathogens that cause hepatitis, diarrhea, or encephalitis in humans, suggesting the presence of new threats to public health. Our study demonstrates the effectiveness of reusing public sequencing data to identify known and unknown viral infections, indicating that future continuous monitoring of public sequencing data by metagenomic analyses would help prepare and mitigate future viral pandemics.

## Introduction

RNA viruses have caused numerous emerging diseases; for example, it was reported that 94 % of zoonoses from 1990 to 2010 were caused by RNA viruses (1). Mammalian and avian species are especially high-risk transmission sources for zoonotic viruses because of their frequent contact with humans as livestock, bushmeat, companion, or laboratory animals (2). Additionally, the spread of viral infectious diseases in livestock animals impacts sustainable food security and economic growth (3). Thus, large-scale surveillance of RNA viral infections in these animals would help monitor infections of known and unknown viruses that can cause outbreaks in humans and domestic animals.

Metagenomic analysis can identify viruses with low bias and has substantially contributed to elucidating virus diversity for more than a decade (4). With the increase in research using metagenomic analysis, new virus species, genera, and families have been successively established by the International Committee on Taxonomy of Viruses (ICTV) (5). However, a previous study estimated the existence of at least 40,000 mammalian viral species (6), which far exceeds the number of viral species classified by the ICTV to date (5, 7). Therefore, further research is needed to understand viral diversity and prepare for future viral pandemics. The amount of RNA-seq data in public databases is growing exponentially (8); however, only limited studies have examined for viral infections in publicly available sequencing data (9–11). The public data are derived from samples with various research backgrounds and may contain a wide variety of viral sequences. Therefore, analyzing publicly available RNA-seq data can be an effective way to assess the spread of viral infections and discover novel viruses.

In this study, we analyzed more than 46,000 RNA-seq data to screen hidden RNA virus infections in mammalian and avian species and identified approximately 900 infections. We also discovered six nearly complete viral genomes in livestock, wild, and laboratory animals. Phylogenetic analyses showed some of the novel viruses were closely related to human pathogenic viruses, suggesting the potential risk of causing disease in humans. Furthermore, the viral infections were identified in several individuals collected by independent studies, indicating that their infections may have already spread in the natural host population. Our findings demonstrate the reusability of public sequencing data for surveying viral infections that may present a threat to public health.

## Results

### Detection of RNA viral infections hidden in public sequence data

To detect RNA viral infections in mammalian and avian RNA-seq data, we first performed *de novo* sequence assembly (**Fig. 1A**). We then performed BLASTX screening using contigs to extract RNA virus-derived sequences. Among 422,615,819 contigs, we identified 17,060 RNA virus-derived sequences. The median length of the viral contigs was 821 bp (**Fig. 1B**), which was shorter than the genomic size of RNA viruses (**Fig. 1C**). These results indicate that most viral contigs were detected as partial sequences of the viral genome, and several contigs may have originated from the same viral infection event. Therefore, we sought to determine the viral infections in each sequencing data by the alignment coverage-based method to avoid double counting (**Fig. 1A** and **details in Materials and Methods**). Briefly, we constructed sequence alignments by TBLASTX using the viral contigs in each RNA-seq data and reference viral genomes, and then calculated the alignment coverage between the viral contigs and each viral reference sequence. Here, we defined a viral infection when the alignment coverage exceeded the threshold (more than 20 %). This threshold was determined using sequencing data obtained from viral infection experiments (**Fig. S1 and details in Materials and Methods**). Finally, we totalized the infections at the virus family level after excluding the viruses inoculated experimentally More than 46,000 mammalian and avian RNA-seq data were used to investigate infections by 33 RNA virus families reported to infect vertebrates. Consequently, we identified 882 infections of 22 RNA virus families in 695 sequencing data from 53 host species (**Fig. 2A**). These results indicate that analyzing public sequencing data by metagenomic analysis is useful for identifying hidden viral infections.

**Figure 1.**
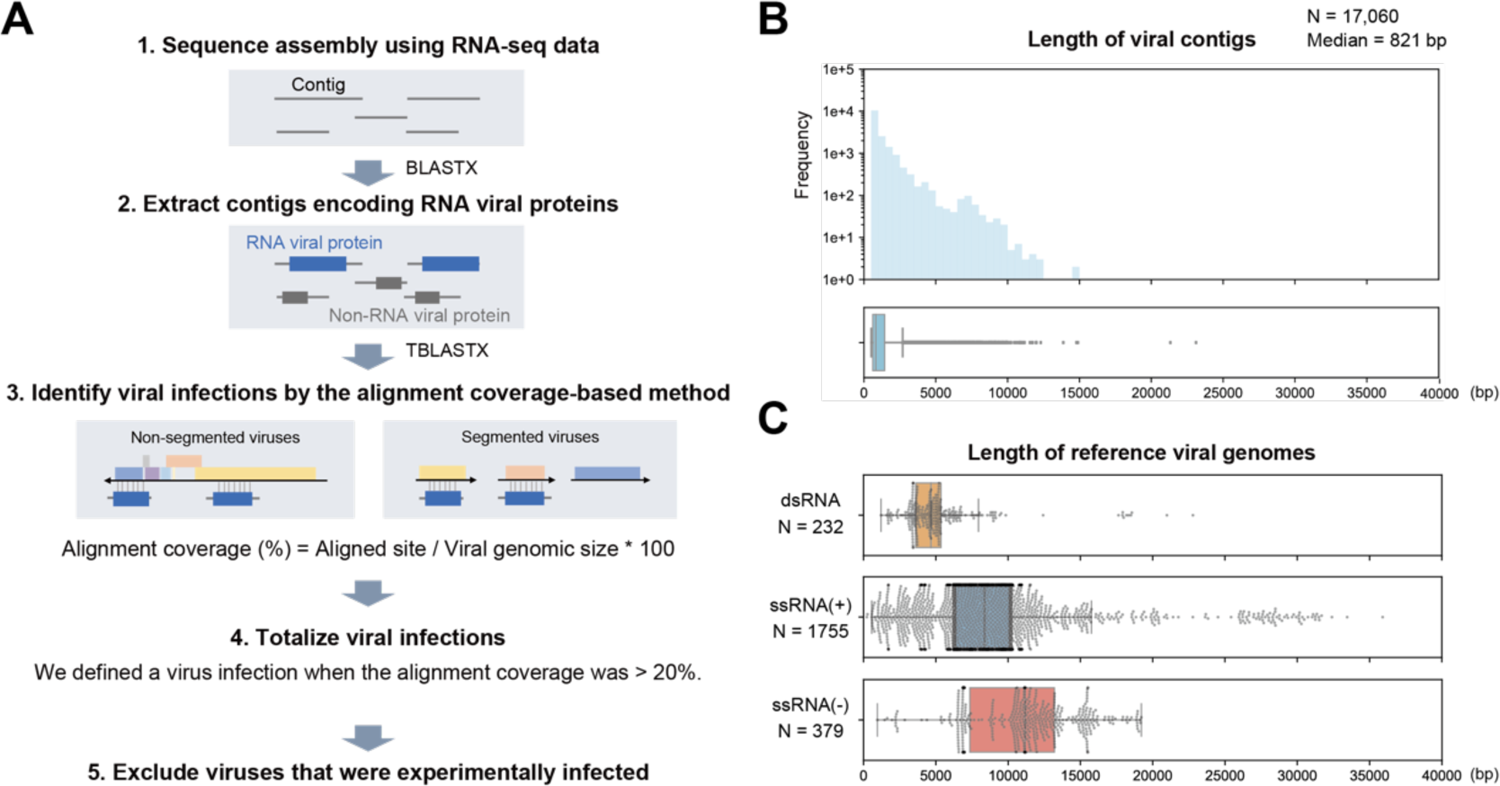
Strategy for detecting viral infections in public RNA-seq data. (A) Schematic diagram of the procedure for detecting viral infections. First, we performed *de novo* sequence assembly using publicly available mammalian and avian RNA-seq data. Next, we extracted contigs encoding RNA viral proteins by BLASTX. Third, we constructed sequence alignments by TBLASTX using the viral contigs in each RNA-seq data and reference viral genomes because most viral contigs were shorter than complete viral genomes, as shown in (B-C). The alignment coverage is defined as the proportion of aligned sites in the entire reference viral genome. Fourth, we determined a viral infection when the alignment coverage was > 20 %. Finally, we totalized the infections at the virus family level after excluding experimentally infected viruses (**details in Materials and Methods).** (B) Distributions of viral contig length: histogram (upper panel) and box plot (lower panel). The x-axis indicates the viral contig length. Among 17,060 viral contigs, the median length was 821 bp. (C) Length of reference viral genomes. Each panel corresponds to the Baltimore classification: the upper, middle, and lower panels show double-stranded RNA (dsRNA) viruses, positive-sense single-stranded RNA (ssRNA(+)) viruses, and negative-sense single-stranded RNA (ssRNA(-)) viruses, respectively. The x-axis indicates the viral genome size. These viral genomes were obtained from the RefSeq genomic viral database. The genomic size of segmented viruses is the sum length of all segments in a virus species.

**Figure 2.**
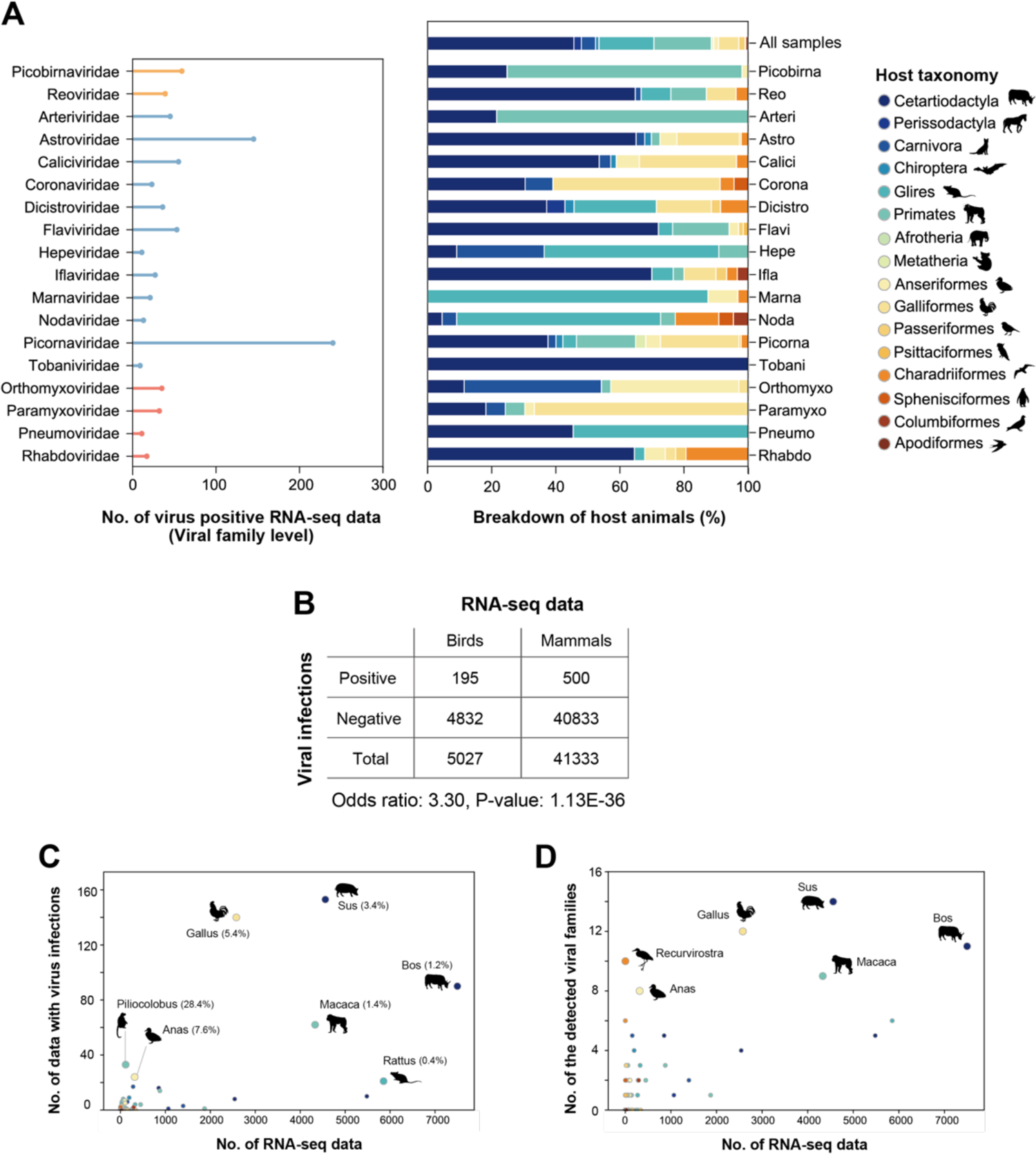
RNA viral infections in the public sequencing data. (A) RNA viral infections detected in public sequencing data. Left panel: the x-axis indicates the number of virus-positive RNA-seq data, and the y-axis indicates viral families. Although infections by 22 RNA viral families were identified in this study, 18 families that were detected in more than five RNA-seq data are shown here. Bar colors correspond to the Baltimore classification, dsRNA viruses (orange), ssRNA(+) viruses (blue), and ssRNA(-) viruses (red). Right panel: breakdown by host animals in which viral family infections were detected. The filled colors correspond to the host taxonomy shown in the legend. The top row indicates the animal-wide breakdown of all RNA-seq data used in this study. (B) Comparison of viral detection rate between avian and mammalian samples. The table shows the number of RNA-seq data with and without viral infections. The odds ratio and p-value were obtained by Fisher’s exact test. (C) Scatter plot between the numbers of RNA-seq data investigated in this study (x-axis) and those with viral infections (y-axis). Each dot indicates the animal genus. Dot colors correspond to the host taxonomy shown in (A). The animal genera, in which viral infections were detected in ≥ 20 samples, are annotated with the representative animal species silhouettes. The percentages in parentheses indicate the ratio of virus-positive RNA-seq data to the investigated data. (D) Scatter plot between the number of RNA-seq data investigated in this study (x-axis) and those of detected viral families (y-axis). Each dot indicates the animal genus. Dot colors correspond to the host taxonomy shown in (A). The animal genera, in which ≥ eight viral families were detected, are annotated with the representative animal species silhouettes.

### Frequent detection of diverse virus families in bird samples

Many viral infectious diseases associated with birds have been reported so far (12), such as influenza A virus (13, 14) and West Nile virus (15). In this study, we frequently detected viral infections in bird samples (**Fig. 2B**). The odds ratio of RNA virus detection in birds compared with that in mammals was 3.3. Furthermore, among the investigated species, we found relatively high viral detection rates in Gallus and Anas species at 5.4 % and 7.6 %, respectively (**Fig. 2C**). We also found infections of 12 and 8 virus families in Gallus and Anas species, respectively (**Fig. 2D**). These results indicate that birds, especially Gallus and Anas species, are frequently infected with various virus families, suggesting that these species are reservoirs for a wide variety of viruses (**see Discussion**).

### Identification of unknown reservoir hosts at virus family levels

To identify novel virus-host relationships at virus family levels, we compared our data with known virus-host relationships provided in the Virus-Host DB (16) (**Fig. 3A**). This database lists virus-host relationships based on the identification of viral sequences from a host animal. We found 50 newly identified virus-host relationships using this database for comparison, and 17 of them were identified with more than 70 % alignment coverage. Notably, we identified nearly complete genomic sequences classified into the family *Hepeviridae* in Spalax and Galago species for the first time. These discoveries expanded our understanding of hepeviral host ranges (details of the viral characteristics are described in the section: “*Hepeviruses in blind mole-rats and a galago: expanding understanding of the hepatitis E virus host range*”). A novel relationship was also identified between the family *Rhabdoviridae* and Recurvirostra species. We did not perform further investigations because the complete rhabdovirus genome could not be obtained, although the alignment coverage was more than 70 %. Additionally, novel virus-host relationships were also found in the families *Dicistroviridae*, *Iflaviridae*, *Marnaviridae*, and *Nodaviridae*, suggesting that these viral host ranges are broader than previously expected. It should be noted that these relationships might be due to contamination from environmental viruses, because few species in these viral families have been reported to infect mammals or birds (17–20) (**see Discussion**).

**Figure 3.**
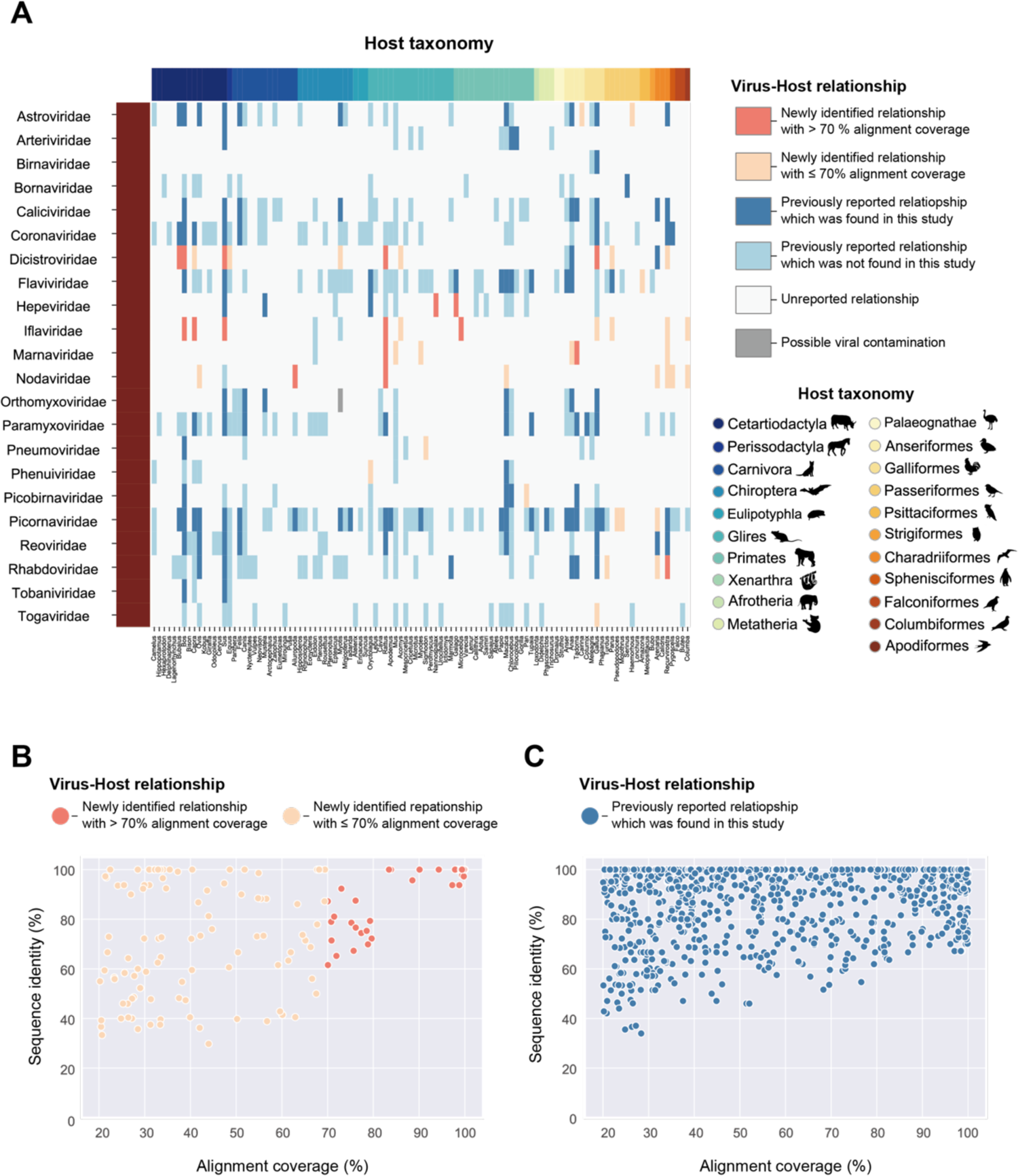
Search for unknown reservoir hosts and novel virus sequences. (A) Heatmap showing the newness of virus-host relationships. Rows indicate viral families that reportedly infect vertebrate hosts. Columns indicate animal genus, and filled colors correspond to the host taxonomy shown in the lower right corner. Heatmap colors are according to six categories of virus-host relationships shown in the upper right corner: a relationship was newly identified in this study, and the viral infection was detected with > 70 % alignment coverage (coral), a relationship was newly identified in this study, but the viral infection was detected with ≤ 70 % alignment coverage (salmon), a relationship was previously reported, and the viral infection was also detected in this study (blue), a relationship was previously reported, but the viral infection was not detected in this study (light blue), a relationship was unreported so far (white), and a relationship was newly identified in this study, but it may be attributed to contamination (gray) (**see Discussion**). (B-C) Scatter plot between alignment coverages (x-axis) and sequence similarities with known viruses (y-axis). Each dot represents the viral infections identified in this study. Viral infections related to novel virus-host relationships are shown in (B), and those related to known relationships are shown in (C). The dot colors correspond to virus-host relationships shown in (A). Sequence identity represents the maximum value of the percentage of identical matches obtained by TBLASTX.

### Investigation of novel viruses with complete genomic sequences

To identify novel sequences comparable to a complete viral genome, we simultaneously analyzed sequence similarity with known viruses and the alignment coverages with reference viral genomes (**Figs. 3B-C**). We found some viral sequences showing low sequence similarity with known viruses and high alignment coverage, which were expected to be novel viruses with a nearly complete genome. Therefore, we further characterized these viral sequences by phylogenetic analyses, annotations of viral genomic features, and quantification of viral reads in RNA-seq data (**Figs. 4-5 and S2-3**). Consequently, we discovered six viruses: hepatovirus in a goat, hepeviruses in blind mole-rats and a galago, astrovirus in macaque monkeys, parechovirus in a cow, and pegivirus in tree shrews.

**Figure 4.**
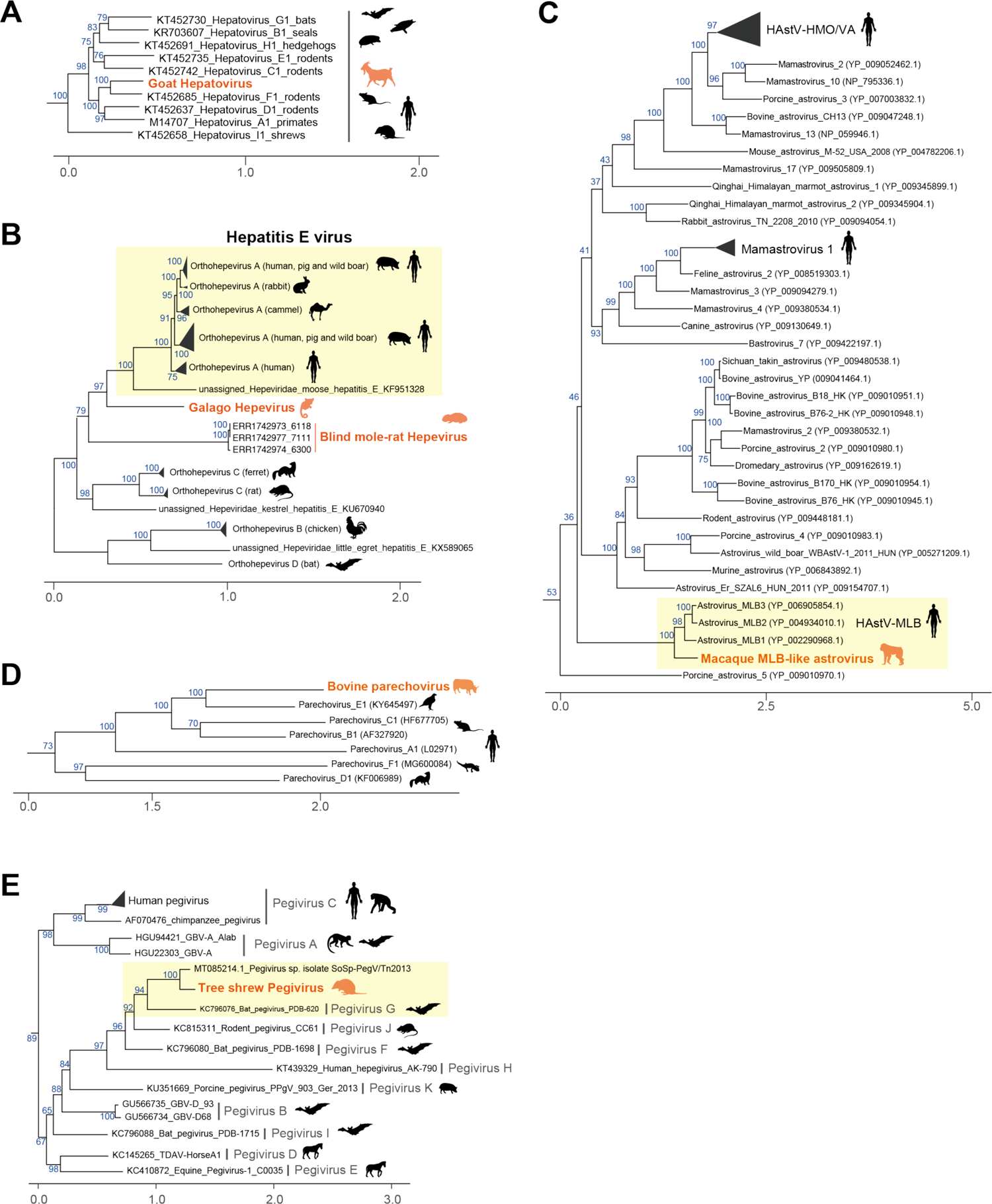
Characterization of virus sequences identified in this study. (A-E) Phylogenetic analyses: the genus *Hepatovirus* of the family *Picornaviridae* (A), the family *Hepeviridae* (B), the genus *Mamastrovirus* of the family *Astroviridae* (C), the genus *Parechovirus* of the family *Picornaviridae* (D), and the genus *Pegivirus* of the family *Flaviviridae* (E). These phylogenetic trees were constructed based on the maximum likelihood method (**details in Materials and Methods**). The orange labels indicate viruses identified in this study, and the colored animal silhouette indicates the viral host species. The black label and animal silhouette indicate known viruses and their representative hosts, respectively. Scale bars indicate the genetic distance (substitutions per site). The blue labels on branches indicate the bootstrap supporting values (%) with 1,000 replicates. Yellow boxes highlight viruses genetically similar to the novel virus identified in this study.

**Figure 5.**
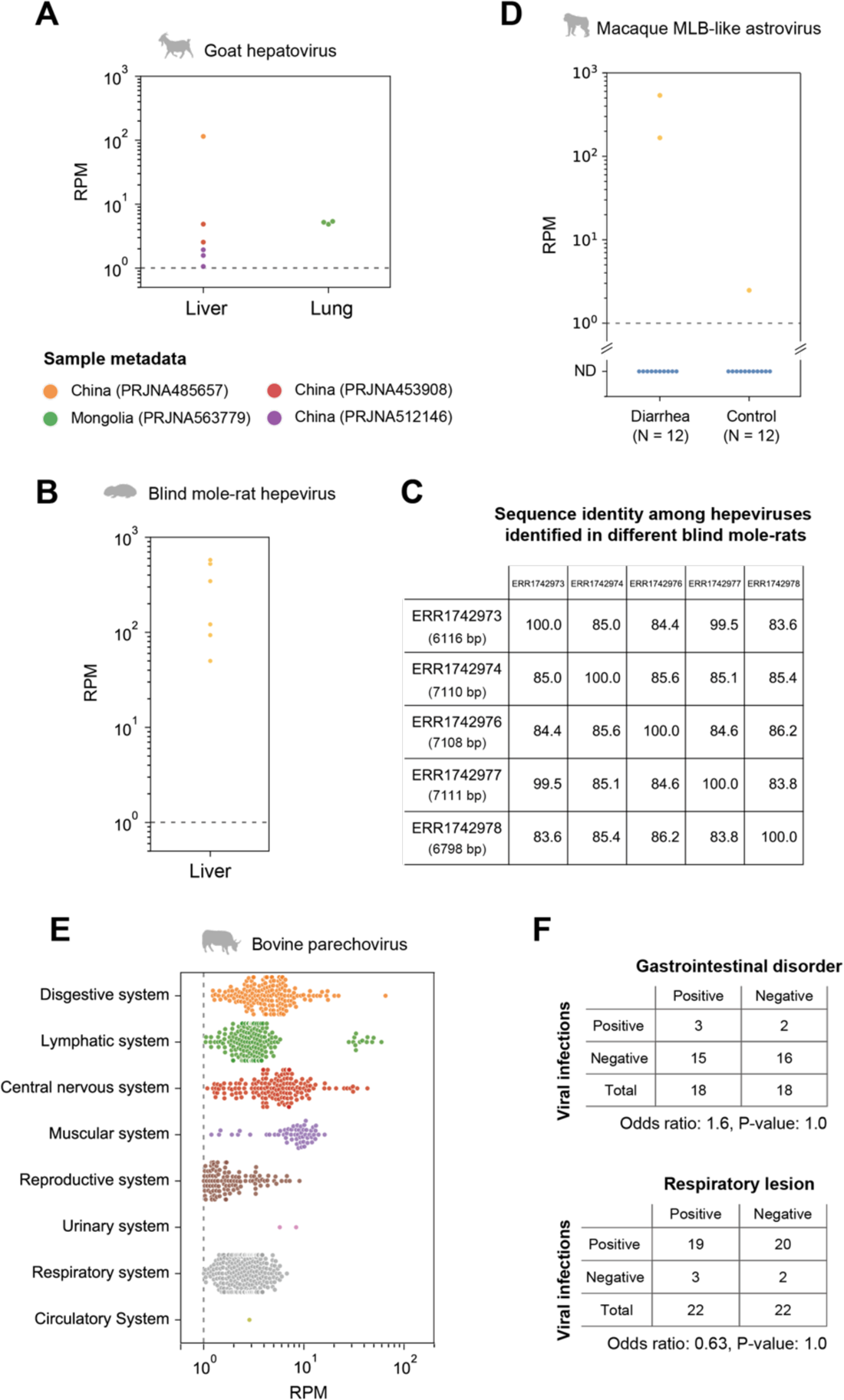
Detection of viral infections in the natural host population. (A, B, and E) Investigation of viral infections in the natural host population by quantifying viral reads: goat hepatovirus (A), blind mole-rat hepevirus (B), and bovine parechovirus (E). Panel indicates the viral read amount (read per million reads [RPM]) in each tissue or organ system. The gray dotted line indicates the criterion used to determine viral infections (RPM: 1.0). The lower panel in (A) represents the sample metadata. (C) Comparison of nucleotide sequence identity among the hepeviral sequences identified in five different blind mole-rats. The numbers in parentheses in the row indicate the total number of aligned sites between the viral contigs identified in each individual and the blind mole-rat hepevirus identified in ERR1742977. (D) Quantification of the macaque MLB-like viral infection levels in the patient with diarrhea and control macaque monkeys. The x-axis indicates the diagnosis for the 24 monkeys, and the y-axis indicates the RPM. The average RPM for each individual is plotted because six samples were collected from each individual. The dotted line indicates the criterion used for detecting viral infections (RPM: 1.0). We considered samples with RPMs below the criterion as non-detectable (ND). (F) Association between the parechovirus infections and symptoms. The tables show the number of RNA-seq data with and without the parechovirus infections in two independent studies, which provide diagnostic information: gastrointestinal disorder (upper panel) and respiratory lesion (lower panel). The odds ratio and p-value were obtained by Fisher’s exact test.

### Goat hepatovirus: the first report on hepatoviral infections in livestock animals

Hepatitis A virus (HAV), belonging to the genus *Hepatovirus* of the family *Picornaviridae*, can cause acute and fulminant hepatitis and is typically transmitted via fecal-oral routes, including contaminated water or foods (21). The World Health Organization (WHO) reported that HAV infections resulted in the death of over 7,000 people in 2016 (https://www.who.int/news-room/fact-sheets/detail/hepatitis-a). Here, we identified a hepatoviral infection in a goat sample (**Fig. 4A**). To our knowledge, this is the first report of hepatoviral infection in livestock animals.

We further analyzed the hepatovirus prevalence in a natural host population by quantifying the viral reads in other goat RNA-seq data because this virus was initially identified in only one goat sample. Among 1,593 samples, we found the viral infections in nine samples from four independent studies with > 1.0 read per million reads (RPM) (**Fig. 5A and Dataset S7**). The goat hepatoviral infections were detected in liver and lung samples, suggesting that the goat hepatovirus can infect tissues other than the liver. Although the lungs are not considered preferential tissues for hepatoviral replication, a previous report also detected hepatoviral RNAs in the seal lungs (22). The infected goat samples were collected in East Asia, including China and Mongolia. Therefore, goat hepatoviruses may be prevalent in the natural host population, suggesting this virus can be a new threat to public health through the contamination of water and foods by infected animals.

### Hepeviruses in blind mole-rats and a galago: expanding understanding of the hepatitis E virus host range

Several million infections of hepatitis E virus (HEV) are estimated to occur worldwide; the WHO reported approximately 44,000 deaths due to HEV infection in 2015 (https://www.who.int/news-room/fact-sheets/detail/hepatitis-e). Here, we found hepeviruses, classified into the same viral family as HEV, in blind mole-rats and a galago for the first time (**Fig. 3A**). Phylogenetic analysis indicated that these hepeviruses formed a single cluster with moose HEV (23) and members of Orthohepevirus A that infect humans, pigs, rabbits, and camels (24) (**Fig. 4B**). However, the hepeviruses identified in this study appeared to have an early divergence from the HEV common ancestor. These results suggest a high diversity and broader host range of HEV-like viruses.

The blind mole-rat hepevirus was identified in the host livers, which coincides with the tissue tropism of HEV (25). Additionally, we found that the 3’-portion of the blind mole-rat hepevirus genome was highly transcribed (**Fig. S3B**), suggesting the transcription of subgenomic RNAs (26). In contrast, we could not determine the tissues infected by the galago hepevirus because the relevant metadata were not available. Further, we did not observe a clear read-mapping pattern that suggests any subgenomic RNA transcription in the galago sample (**Fig. S3C**).

We also investigated the spread of these hepeviruses in a natural population using RNA-seq data from blind mole-rats and galagos. Among 91 RNA-seq data from blind mole-rats, we detected the hepeviral infections in six samples (**Fig. 5B**). These infected individuals were captured and kept as laboratory animals in Israel by the same research (**Dataset S8**). There were two possibilities about when the hepeviruses have infected blind mole-rats: the hepeviruses had already infected these blind mole-rats when they were captured, or the viral infections had spread during the maintenance of these individuals in the laboratory. To explore these possibilities, we investigated the inter-individual diversity of the hepevirus sequences and found that these blind mole-rats were infected with relatively diverse hepeviruses representing nucleotide sequence identities ranging from 83.6 % to 99.5 % (**Fig. 5C**). These results suggested that several individuals had already been infected with distinct hepeviruses in the wild before being captured. The galago hepeviral infections were detected in only two samples originating from a study in which we first identified the virus (**Dataset S9**). This may be simply because only four galago RNA-seq data obtained from the same study were available. Taken together, we suggest that these hepeviruses can become a new threat to public health, similar to HEV.

### MLB-like astrovirus detected in macaque monkeys with chronic diarrhea

We found astroviruses genetically similar to human astrovirus MLB (HAstV-MLB) in fecal samples of macaque monkeys (**Fig. 4C**). Although HAstV-MLB infections are typically asymptomatic (27, 28), several studies have reported the viral detection in cases with diarrhea (29), encephalitis (30), or meningitis (31). Interestingly, the macaque MLB-like astrovirus was found in macaque monkeys with chronic diarrhea. We analyzed the viral read amounts in the patient (n = 12) and control (n = 12) monkeys to assess the association between MLB-like astroviral infections and symptom prevalence (**Fig. 5D and Dataset S10**). Abundant MLB-like astroviral reads were detected in two patients, suggesting that the viral infections are associated with host symptoms. However, we did not observe the viral infection in other patients; further, we found the infection in a control individual, although the viral read amount was approximately 100 times less than those of the patients. Additionally, a previous study reported that monkeys, in which partial sequences of MLB-like astroviruses were detected, had no obvious clinical signs, including diarrhea (32). Thus, further experiments are needed to clarify the pathogenesis of MLB-like astrovirus. Considering that there is no current experimental system for examining HAstV-MLB infections (28), our findings suggest that macaque monkeys can be used as animal model systems for researching MLB-like astroviruses.

### Silent infections of bovine parechovirus having a broad tissue tropism

Human parechovirus infection is especially problematic in infants and young children. Although most parechovirus infections are considered asymptomatic, their infections have been reported in patients with respiratory, digestive, and central nervous system disorders (33). In this study, we identified a parechovirus, classified into the family *Picornaviridae*, in the lower digestive tract of a cow (**Fig. 4D**). Despite the broad host range of parechovirus, including mammals, birds, and reptiles (34), to our knowledge, this is the first report on parechovirus infections in livestock animals.

Phylogenetic analysis indicated that this parechovirus was closely related to the falcon parechovirus, a member of Parechovirus E. Next, we compared the bovine parechovirus with the ICTV species demarcation criteria (34) to investigate whether this virus is a novel species (**Fig. S2B**). Consequently, we found that the bovine parechovirus was distant enough from other known parechovirus species and could be considered a separate species based on the following criteria: divergence of amino acid sequences in polyprotein (37.8 %), P1 protein (37.8 %), and 2C+3CD (29.9 %) protein. Therefore, we propose that this virus belongs to a new species in the genus *Parechovirus*.

We also investigated the prevalence of this parechovirus infection in a natural host population using public cow RNA-seq data (**Fig. 5E and Dataset S11**). Among 8,284 samples, we detected the parechovirus infections in 944 samples from eight independent studies with > 1.0 RPM. The viral infections were detected in various tissues, such as the digestive, lymphatic, and central nervous systems. These results suggest a broad tissue tropism of the bovine parechovirus. To assess the parechovirus pathogenicity, we analyzed the viral prevalence among 36 or 44 samples with a diagnosis for a gastrointestinal disorder or respiratory lesion, respectively. We did not observe a significant association between the viral infections and the presence/absence of symptoms in these two studies (**Fig. 5F**). These results indicate that bovine parechovirus infections may be asymptomatic, similar to the typical outcome of human parechoviral infections. Furthermore, this also suggests that infected cows can spread parechoviral infections as silent reservoirs.

### Geographical expansion of tree shrew pegivirus infection associated with host migration

We found a pegivirus belonging to the genus *Pegivirus* of the family *Flaviviridae* in tree shrew liver samples. Phylogenetic analysis indicated that this pegivirus was closely related to the Pegivirus G identified in various bat species (**Fig. 4E**). According to the ICTV species demarcation criteria (35), this virus appeared to be the same species as Pegivirus G because the amino acid sequence identity in the NS5B gene was 70.9 % (**Fig. S2C**). These results indicate that Pegivirus G can infect distinct host lineages: tree shrews and bats.

We also investigated the pegiviral infections in other tree shrew samples by read mapping analysis. Among the 59 samples, the pegiviral infections were detected in four samples collected from a research colony in the United Kingdom (**Dataset S12**). A recent report partially identified a pegiviral sequence (GenBank: MT085214.1) in tree shrews collected in Southeast Asia (36), which showed 84.9 % nucleotide sequence identity to the pegivirus identified in this study (**Fig. 4E**). These results indicate that tree shrew pegivirus infections were found in both Asia and Europe, suggesting an expanding geographic distribution of Pegivirus G along with host animal transportation as experimental resources. Thus, the global trade of host animals may lead to spreading pegiviral infections hidden in tree shrews.

## Discussion

Metagenomic analysis is a powerful approach for surveying viral infections (4, 5). Extensive deep sequencing data have been accumulated in public databases, which can be used for identifying viral infections. In this study, we analyzed the publicly available RNA-seq data to search for hidden RNA viral infections in mammalian and avian species and subsequently identified approximately 900 infections by 22 RNA virus families (**Figs. 1 and 2**). These results indicate that reusing public sequencing data is a cost-effective approach for identifying viral infections. Furthermore, we discovered six novel viral genomes in livestock, wild, and experimental animals (**Fig. 4**). Some of these viruses were detected in different individuals, suggesting that the viral infections may have already spread in the natural host population (**Fig. 5**). Overall, our work demonstrates the reusability of public sequencing data for surveying infections by both known and unknown viruses.

In this study, we determined viral infections by a combination of sequence assembly and the alignment coverage-based method to solve several issues in viral metagenomic analysis (**Fig. 1A**). One of the problems is detecting infections in data with a small number of viral reads because almost all public sequencing data were collected without using virus enrichment strategies. The result that most virus contigs were shorter than the reference viral genomes reflects this difficulty (**Figs. 1B-C**). To resolve this issue, we determined viral infections by the alignment coverage-based method, which uses relatively short viral sequences as clues (**Figs. 1A and S1**). Consequently, we succeeded in detecting approximately 900 RNA viral infections in public sequencing data (**Fig. 2A**). Another problem in viral metagenomic analysis is the viral detectability depending on sequence similarity with known viruses. We here discovered six nearly complete viral genomes (**Fig. 4**) by sequence assembly and BLAST screening (**Fig. 1A**). Notably, these viral infections were undetectable in almost all samples, even at the virus family and genus levels, by the NCBI SRA Taxonomy Analysis Tool (https://github.com/ncbi/ngs-tools/tree/tax/tools/tax), which determines the taxonomic composition of reads in the RNA-seq data without sequence assembly (**Dataset S7-S12**). These results indicate that our method can identify novel viruses with full-length genomes, which would effectively elucidate virus diversity. Taken together, our strategy using sequence assembly and the alignment coverage-based method can efficiently detect known and unknown viral infections in publicly available sequencing data.

However, there are still several challenges for identifying viral infections in public sequencing data. First, we could not determine complete viral sequences mostly (**Figs. 3B and 3C**). Further improvement in sequence assembly efficiency (37) or integrative analysis using short- and long-read sequence datasets (38) can solve this problem. Second, there may be a bias in virus detection using public sequencing data depending on their genomic types. Among the 882 viral infections identified in this study, 77.0% were positive-sense single-stranded RNA (ssRNA(+)) viral infections, whereas 11.5% and 11.5% were double-stranded RNA and negative-sense single-stranded RNA viral infections (**Fig. 2A**). The RNA-seq procedure, such as enrichment of polyadenylated (poly-A) transcripts, can be relevant to this bias because many ssRNA(+) viruses have a poly-A tract at the 3’-end of their genome (39). Alternatively, this bias may result from a repertoire of reference viral genomes used for the viral search (**Fig. 1C**), which can be solved in the future by database expansion. Thirdly, our method demands relatively abundant computational resources, including operation time, for determining viral infections in each RNA-seq data. We reconstructed viral sequences from RNA-seq data according to several steps: mapping analysis for excluding host transcripts, de novo sequence assembly using unmapped reads, and BLASTX screening for identifying viral sequences (**Fig. 1A**). In contrast, another study performed a search for viral RNA-dependent RNA polymerase proteins in translated nucleotide sequences, which enabled the authors to screen for viral infections in approximately 5.7 million public sequencing data within 11 days (11). Considering that the number of public sequencing data will keep increasing, platform development and maintenance, which can save computational resources, would be necessary for continuing such viral surveillance.

Another challenge in viral metagenomic analysis using public data is distinguishing true viral infections from contamination. To address this issue, we performed integrative analyses using sample metadata and sequence information, including sequence similarity and alignment coverage with known viruses (**details in Materials and Methods**). Consequently, we found several possible contamination cases: influenza A virus in a Myotis bat, vesicular stomatitis Indiana virus (VSV) in cultured chicken cells, mammalian rubulavirus 5 (PIV5) in cultured cells and quail egg samples, and Kadipiro virus (KDV) in rat samples (**Fig. 3A and Dataset S3**). For example, the influenza A viral nucleotide sequence identified in a bat sample showed 100 % similarity to a laboratory strain of influenza A virus (A/WSN/1933(H1N1)). Considering that the bat sample was collected in 2012, it is difficult to expect that such a highly similar influenza A virus was maintained for approximately 80 years. Likewise, the infections of VSVs and PIVs were also identified with approximately 100 % sequence similarity to the reference viral sequences (**Dataset S3**). VSV is frequently used as experimental tools; for example, as a pseudotype virus (40). Previous studies also have reported possible contamination of PIV5 in cultured cells (41, 42). Additionally, it was reported that KDV RNA might be a contaminant in the nucleic acid extraction kit (43). Therefore, we excluded these viral infections to avoid counting false positives. These cases emphasize the importance of multilayered validations for viral infections found by viral metagenomic analysis alone.

Further research efforts to elucidate viral diversity are necessary to prepare for a possible future viral pandemic (1, 5). A strategic approach, such as determining the host samples used for virus search based on the expectation of viral infection frequency or viral diversity, would be necessary. It has been discussed that birds may be high-risk viral hosts of zoonoses because of their high species diversity and wide habitat range (12). In this study, we found that viral infections were more frequently detected in birds, especially Gallus and Anas species (**Figs. 2B-D**). Furthermore, among 217 viral infections identified in Gallus and Anas samples, 78 infections (35.9 %) showed less than 95 % amino acid sequence similarity with known viruses, suggesting that these sequences may be derived from unknown viruses. Therefore, further viral metagenomic analyses targeting bird samples may effectively detect viral infections, including unknown ones.

In conclusion, we demonstrated the reusability of public sequencing data for monitoring viral infections and discovering novel viral sequences, and elucidated diverse RNA viruses hidden in animal samples. Our findings also emphasize the necessity of continuous surveillance for viral infections using public sequencing data to prepare for future viral pandemics, as well as the importance of developing a fundamental bioinformatics platform for surveillance (11, 44).

## Materials and Methods

### Sequence assembly using publicly available RNA-seq data

We collected RNA-seq data of 41,332 mammals (169 genera and 228 species) and 5,027 birds (70 genera and 83 species) from the NCBI Sequence Read Archive (SRA) database (8) during June and July 2019 according to the following search conditions: [(“Mammalia”[Organism] OR “Mammals”[All Fields]) AND (“biomol rna”[Properties] AND “library layout paired”[Properties] AND “filetype fastq”[Properties]) NOT (“Homo sapiens”[Organism]) NOT (“Mus musculus”[Organism])] and [(“Aves”[Organism] OR “Aves”[All Fields]) AND (“biomol rna”[Properties] AND “library layout paired”[Properties] AND “filetype fastq”[Properties])]. The RNA-seq data were downloaded from the NCBI SRA database by pfastq-dump (https://github.com/inutano/pfastq-dump) and preprocessed using fastp (version 0.20.0) (45) with options “-l 35”, “-y-3”, “-W 3”, “-M 15”, and “-x”.

Sequence assembly was conducted by 1) mapping reads to the host or sister species genome and 2) *de novo* assembly of sequences using unmapped reads. First, we performed a mapping analysis to exclude the reads originating from host transcripts and endogenous viral elements. We mapped the reads in each RNA-seq data to the host genome by HISAT2 (version 2.1.0) (46) with the default parameters or used the sister species genomes of the host in the same genus when the host genome data were not available. Unmapped reads were extracted by Samtools (version 1.9) (47) and Picard (version 2.20.4) (http://broadinstitute.github.io/picard). When the relevant genome data were unavailable, the preprocessed reads were directly used for sequence assembly. Sequence assembly was conducted by SPAdes (version 3.13.0) (48) or metaSPAdes (version 3.13.0) (49) with *k*-mers of 21, 33, 55, 77, and 99. Finally, we excluded contigs with lengths shorter than 500 bp by Seqkit (version 0.9.0) (50) and then clustered the contigs showing 95.0 % nucleotide sequence similarity by cd-hit-est (version 4.8.1) (51). Consequently, we obtained 422,615,819 contigs and used them for subsequent analyses. We listed the SRA Run accession numbers, genome files used for mapping analysis, and sequence assembly tools in **Dataset S1**.

### Identification of contigs originating from RNA viruses

To determine the origins of the contigs, we analyzed the sequence similarity between the contigs and known sequences in BLASTX screening (version 2.9.0) (52). First, we performed BLASTX searches with the options “-word_size 2”, “-evalue 1E-3”, and “max_target_seqs 1” using a custom database consisting of RNA viral proteins. We constructed the custom database by downloading the viral protein sequences of the realm *Riboviria* from the NCBI GenBank (version: 20190102) (53) and clustering the sequences showing 98.0 % similarity by cd-hit (version 4.8.1). Second, to confirm that the contigs are not derived from organisms other than viruses, we further performed BLASTX searches with the options “-word_size 2”, “-evalue 1E-4”, and “-max_target_seqs 10” using the NCBI nr database (versions: 20190825-20190909 were used for screening contigs in mammalian data and versions: 20190330-20190403 were used for screening contigs in avian data). We determined the contig origins by comparing the bitscores in the first and second BLASTX screening. Consequently, we obtained 17,060 contigs that were deduced to encode RNA viral proteins.

### Totalization of RNA viral infections in public RNA-seq data

Since most viral contigs were shorter than the reference viral genomic sizes (**Figs. 1B-C**), we sought to determine viral infections based on the alignment coverage-based method (**Fig. 1A**). First, we performed sequence alignment by TBLASTX (version 2.9.0) using viral contigs from the RNA-seq data and complete viral genomes in the NCBI RefSeq genomic viral database (version 20200824). Next, we calculated the alignment coverage with the genome of each viral species: the proportion of aligned sites in the entire reference viral genome. In this study, we considered that an infection of the viral family is present if the alignment coverage was greater than 20 %. Validation of this totalization method and evaluation of the criteria are described in the next section (**Fig. S1**). Furthermore, we manually checked sequences with more than 70 % alignment coverage and more than 95 % identity with known viruses in the TBLASTX alignment to examine possible contamination with laboratory viral strains, as well as experimentally inoculated viruses. We excluded experimentally inoculated viral infections (**Dataset S2**) and possible contamination (**Dataset S3**) from the final totalization (**Fig. 2A**).

### Validation of the procedure used to totalize viral infections

Using samples obtained from viral infection experiments, we first compared the alignment coverage-based method with that based on viral read amounts in order to validate the detection rate of viral infections of our method (**Fig. S1 and Dataset S2**). We obtained the read amounts derived from experimentally infected viruses from the NCBI SRA Taxonomy Analysis Tool results (https://github.com/ncbi/ngs-tools/tree/tax/tools/tax). The calculation procedure for alignment coverage between viral contigs in each RNA-seq data and viral reference genomes is described in the previous section. We observed a positive correlation between the alignment coverage and viral read amounts (Pearson’s correlation coefficient: 0.19, p-value: 1.87E-6) (**Fig. S1A**). Among the samples collected from experiments of viral infections, the true-positive rate (the detection rate of experimentally inoculated viruses) was 88.3 %, and the false-positive rate (the rate that mock samples were determined to be infected samples) was 62.5 % when we used 20 % alignment coverage as the criterion for determining viral infections (**Fig. S1B**). The relatively high false-positive rate may be due to similar amounts of viral reads in some mock samples as those in infected samples (**Fig. S1A**). Next, we analyzed the association between alignment coverages and viral genome size (**Fig. S1C**) because the detectability of viral infections in our method may depend on the reference viral genome size. As expected, we observed a tendency for viruses with small genomes to be detected with relatively high alignment coverage. However, more than 80 % of experimentally infected viral infections were detected with more than 20 % alignment coverage, regardless of the viral genome size. Based on these results, we established the alignment coverage of 20 % to totalize the viral infections. Consequently, we identified a total of 1,410 RNA viral infections, including 503 infections in samples from viral infectious experiments (**Fig. S1D**).

### Collection of information on experimentally infected viruses

To exclude experimentally infected viruses from the final totalization, we investigated the experimental background of RNA-seq data. We first collected the experimental descriptions of RNA-seq data: title and abstract from the NCBI BioProject database (54). Then, we manually checked the terms relevant to viral infections in the descriptions, focusing on viral name abbreviations and viral vector usage. We listed the obtained information about viral infection experiments in **Dataset S2**.

### Summarization of virus-host relationships

To identify novel reservoir hosts at the viral family levels, we compared the virus-host relationships identified in this study with the dataset provided by the Virus-Host DB (version: 20200629) (16). We defined a “novel virus-host relationship” as one in which the viral sequence has not been reported in the host. The virus-host relationships at the viral family level were categorized as 1) a novel relationship detected with > 70 % alignment coverage, 2) a novel relationship detected with ≤ 70 % alignment coverage, 3) a known relationship that was also detected in this study, 4) a known relationship that was not identified in this study, 5) a relationship unreported so far, and 6) a novel relationship, which was possibly derived from contamination (**see Discussion**). To avoid misclassification of the relationships, we analyzed reports manually by searching the NCBI PubMed and Nucleotide databases using the combination of the host genus and viral family names: for example, [“Pan” AND “Picobirnaviridae”]. The results of the manual curation are listed in **Dataset S4**.

### Characterization of viral genomic architectures

Open reading frames (ORFs) and polyadenylation signals in the viral genomes were predicted by the SnapGene software (snapgene.com). The positions of mature proteins, frameshift signal sequences, and subgenomic RNA promoter sequences were predicted based on sequence alignment using novel and reference viral sequences. The sequence alignments were constructed by MAFFT (version 7.407) (55) with the option “--auto”. The reference viral sequences used for the genome annotations are listed in **Dataset S5**. The viral sequences identified in this study are registered under the following accession numbers: BR001715-BR001732 and BR001751.

### Phylogenetic analyses

Multiple sequence alignments (MSAs) of picornaviral P1 nucleotide sequences for **Fig. 4A**, hepeviral ORF1 amino acid sequences for **Fig. 4B**, picornaviral 3D nucleotide sequences for **Fig. 4D**, and flaviviral NS5 nucleotide sequences for **Fig. 4E** were obtained from the ICTV resources (the family of *Picornaviridae*: https://talk.ictvonline.org/ictv-reports/ictv_online_report/positive-sense-rna-viruses/picornavirales/w/picornaviridae/714/resources-picornaviridae, the family of *Hepeviridae*: https://talk.ictvonline.org/ictv-reports/ictv_online_report/positive-sense-rna-viruses/w/hepeviridae/731/resources-hepeviridae, and the family of *Flaviviridae*: https://talk.ictvonline.org/ictv-reports/ictv_online_report/positive-sense-rna-viruses/w/flaviviridae/371/resources-flaviviridae). Because an astroviral MSA was not available in the ICTV resources, we extracted astroviral ORF2 amino acid sequences from RefSeq protein viral database (version 20201007). The MSAs of reference and novel viral sequences were constructed by MAFFT with options “--add” and “--keeplength”. The astroviral MSA was trimmed by excluding sites where > 20 % of the sequences were gaps and subsequently removing sequences with less than 80 % of the total alignment sites. Phylogenetic trees were constructed by the Maximum likelihood method using IQTREE (version 1.6.12) (56). The substitution models were selected based on the Bayesian information criterion provided by ModelFinder (57): GTR+R8 for **Fig. 4A**, LG+F+R4 for **Fig. 4B**, LG+F+R5 for **Fig. 4C**, TVM+R9 for **Fig.4D**, and GTR+R7 for **Fig. 4E**. The branch supportive values were measured as the ultrafast bootstrap by UFBoot2 (58) with 1,000 replicates. Tree visualization was performed by the ggtree package (version 2.2.1) (59). Sequence accession numbers used for the phylogenetic analyses are listed in **Dataset S5**.

### Comparison with the ICTV species demarcation criteria

To assess whether the viruses identified in this study could be assigned to a novel species, we compared their genetic distance with known viruses according to the ICTV species demarcation criteria (34, 35) (**Fig. S2**). Amino acid sequences of the P1 and 3CD genes in hepatoviruses and parechoviruses were extracted by referring to Hepatovirus A (GenBank: M14707.1) and Parechovirus A (GenBank: S45208.1), respectively. Amino acid sequences of the NS3 and NS5B genes in pegiviruses were extracted by referring to Pegivirus A (GenBank: U22303.1). We constructed MSAs using these reference and novel viral sequences by MAFFT with the option “--auto”. We did not analyze other viruses identified in this study because the ICTV did not provide criteria based on the genetic distance. The sequence accession numbers used for these analyses are listed in **Dataset S5.**

### Mapping analyses using viral genomes identified in this study

To verify the quality of sequence assembly, we mapped the reads in the RNA-seq data, in which a novel viral sequence was identified, to the viral genomes by STAR (version 2.7.6a) (60) (**Fig. S3**). The genome indexes were generated with the option “--genomeSAindexNbases” according to each viral genomic size, and mapping analysis was conducted with the options “--chimSegmentMin 20”. The number of mapped reads in each position was counted by Bedtools genomecov (version 2.27.1) (61) with the options “-d” and “-split”.

To identify novel viral infections in other individuals, we analyzed the publicly available RNA-seq data of the host animals by quantifying viral reads (**Figs. 5A, B, and 5E**). We investigated 1,593 goat, 91 blind mole-rat, four galago, 8,282 cow, and 59 tree shrew data for infections of goat hepatovirus, blind mole hepevirus, galago hepevirus, bovine parechovirus, and tree shrew pegivirus, respectively. Mapping analyses were performed using STAR (version 2.7.6a) as described above. The number of total and mapped reads was extracted by Samtools (version 1.5). We considered that there was a viral infection in the sample if the RPM was > 1.0.

We compared the viral read amounts between the patient and control monkeys to investigate the association between chronic diarrhea and MLB-like astrovirus infection (**Fig. 5D**). Viral read amounts were quantified as described above. The average RPM for each individual is plotted in **Fig. 5D** because six samples were collected from each individual. **Dataset S6** shows the SRA Run accession number used to investigate novel viral infections. **Datasets S7-S12** list sample metadata in which the novel viral infections were detected.

### Comparison of hepeviral sequences identified in different blind mole-rats

We compared nucleotide sequence identities among the hepeviral sequences found in five different individuals to predict when these viruses infected the blind mole-rats. The sequence comparison was performed by BLASTN (version 2.11.0) with default parameters. Because most hepeviral sequences were detected as short contigs, sequence identities were represented by the percentage of identical matches in the longest aligned region between the hepeviral sequences (**Fig. 5C**). We also analyzed the total of aligned length between contigs identified in each individual and the hepeviral genome identified in ERR1742977 and confirmed that these contigs covered 86.0-99.9% of the blind mole-rat hepevirus genome.

## Data Availability

Bioinformatics tools and their versions are listed in **Dataset S13**.

## Acknowledgments

We thank Jumpei Ito (Institute of Medical Science, The University of Tokyo, Japan) and Dr. Keiko Takemoto (Institute for Virus Research, Kyoto University, Japan) for their technical support. We are grateful to Dr. Nicholas F Parrish (RIKEN Hakubi Research Team, RIKEN Cluster for Pioneering Research, Yokohama, Japan), Bea Clarise Garcia, Yahiro Mukai, Hsien Hen Lin, and Koichi Kitao (Institute for Frontier Life and Medical Sciences, Kyoto University) for helpful discussions. We thank Editage (http://www.editage.com) for editing and reviewing this manuscript for English language. This study was supported by JSPS KAKENHI JP19J22241 (JK), JP18K19443 (MH); MEXT KAKENHI JP17H05821 (MH) and JP19H04833 (MH); Hakubi project at Kyoto University (MH). Computations were partially performed on the supercomputing systems: SHIROKANE (Human Genome Center, the Institute of Medical Science, The University of Tokyo) and the NIG supercomputer (ROIS National Institute of Genetics).

## Author contributions

MH and JK conceived the study; JK and MH mainly performed bioinformatics analyses; SK supported bioinformatics analyses; JK and MH prepared the figures and wrote the initial draft of the manuscript; all authors designed the study, interpreted data, revised the paper, and approved the final manuscript.

---

**Supplemental Figure 1.**
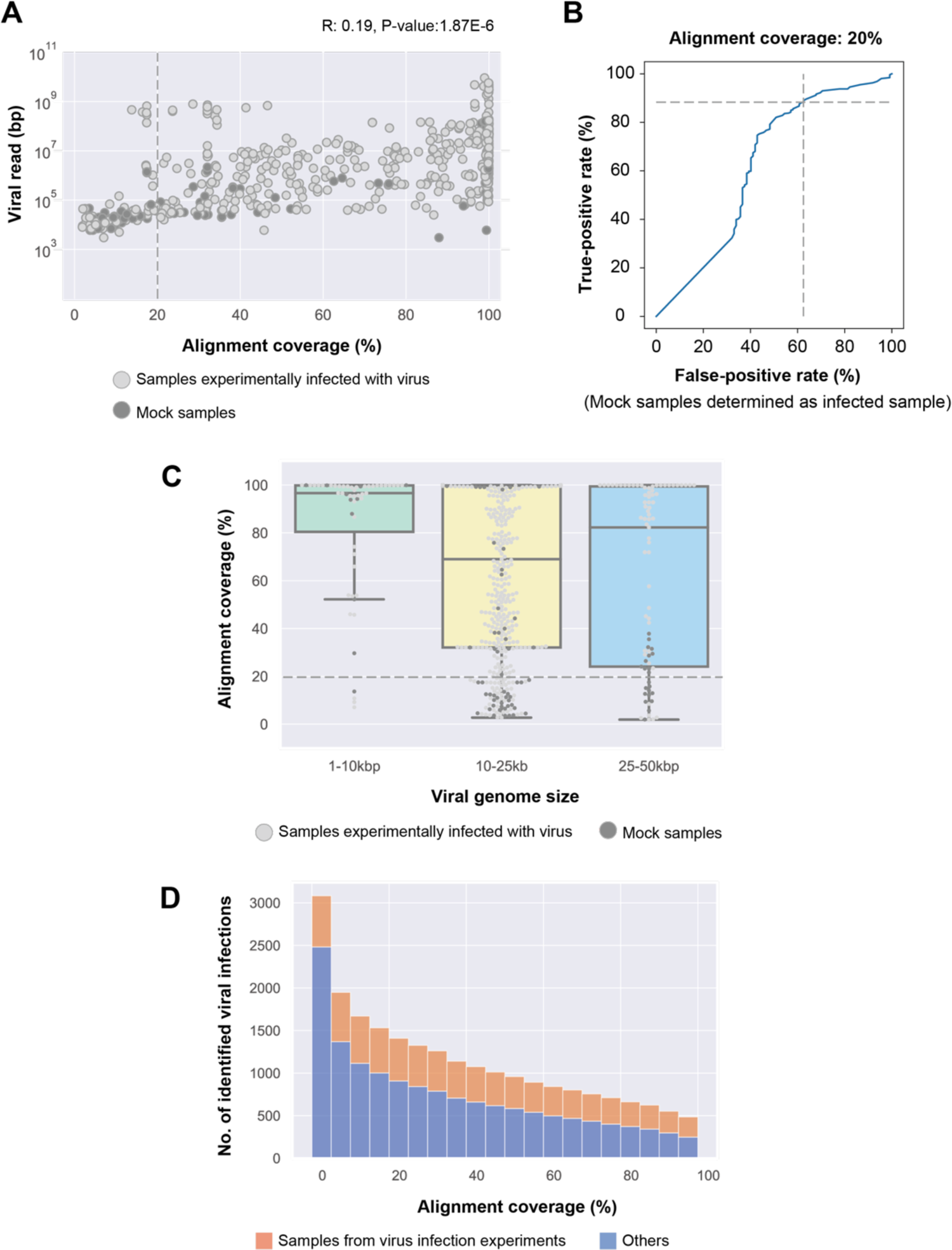
Validation of the alignment coverage-based method for detecting viral infections. (A) Comparison between the alignment coverage-based method and the viral read-based method using samples obtained from viral infection experiments. The x-axis indicates alignment coverage between viral contigs in each RNA-seq data and the reference viral genome used for the experiments. The y-axis indicates the total read length of the virus family used for the experiment, which was obtained from the NCBI SRA Taxonomy Analysis Tool. Light gray dots indicate samples experimentally infected with viruses, and dark gray dots indicate mock samples. R: Pearson’s correlation coefficient. Dotted line indicates 20 % alignment coverage. (B) Changes in the true-positive and the false-positive rates depending on the criteria to determine viral infections. The true-positive rate (y-axis) indicates the number of samples experimentally infected with viruses correctly determined as the infected sample, and the false-positive rate (x-axis) indicates the number of mock samples determined as the infected sample. Dotted line indicates the true-positive rate (88.3 %) and the false-positive rate (62.5 %) when 20 % alignment coverage was used as the criterion (**details in Materials and Methods).** (C) Detection rate of viral infections depending on the viral genome size. Box plots show the distributions of alignment coverage of the viral genome with 1-10kbp (green), 10-25kbp (yellow), and 25-50kbp (blue). Light gray dots indicate samples infected with viruses experimentally, and dark gray dots indicate mock samples. Dotted line indicates alignment coverage. (D) The number of detected viral infections depending on the alignment coverage criteria. The x-axis indicates alignment coverage used as a criterion for defining viral infections. Bar graphs show the number of detected viral infections using the criterion shown on the x-axis. Filled colors indicate infections in samples from viral infection experiments (orange) or those in others (blue). When we used 20 % alignment coverage as the criterion, a total of 1,410 viral infections were identified, including 503 experimentally infected samples.

**Supplemental Figure 2.**
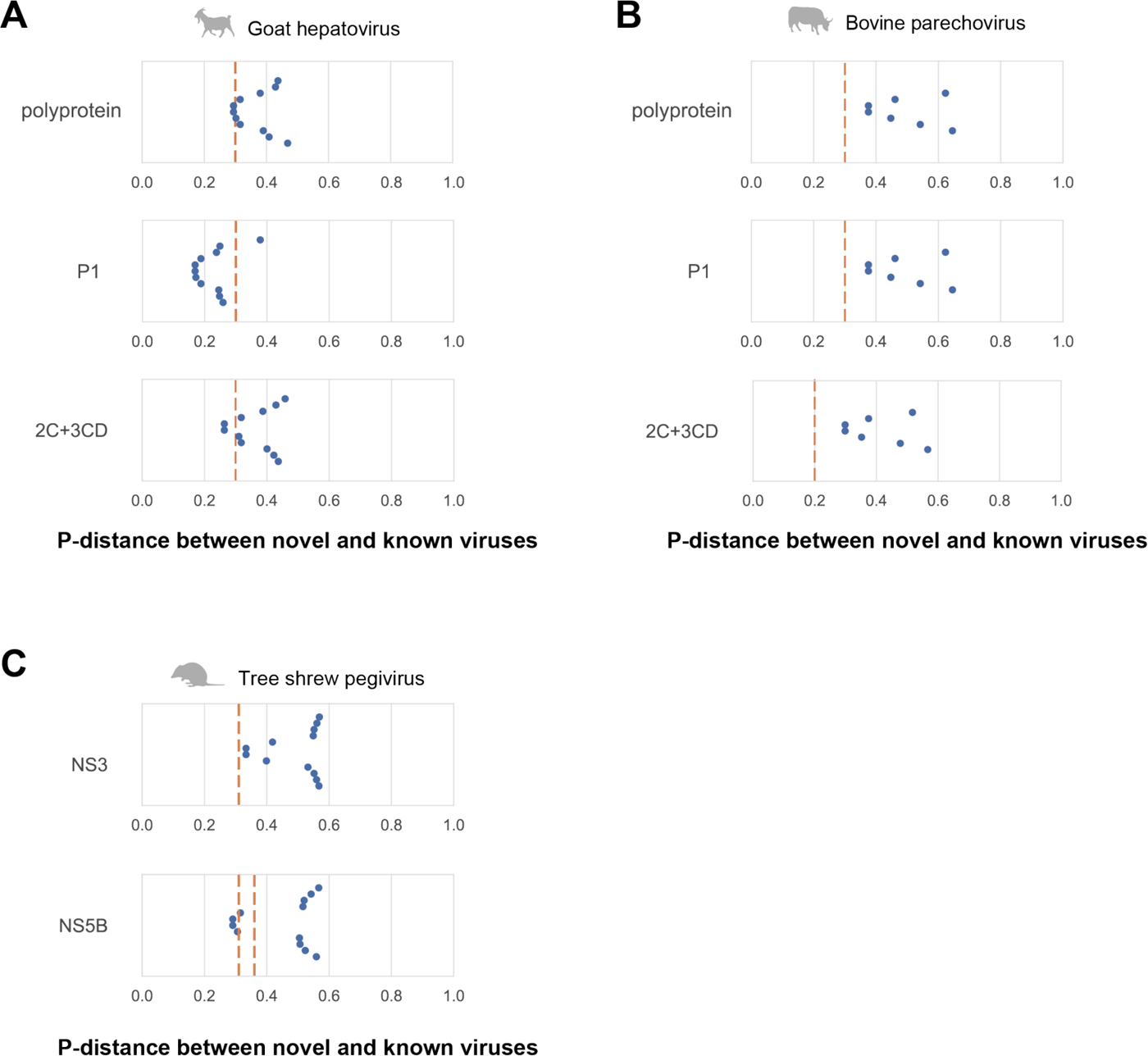
Comparison with the ICTV species demarcation criteria. (A-C) Genetic distance among the amino acid sequences of novel and known viruses in the genera *Hepatovirus* (A), *Parechovirus* (B), and *Pegivirus* (C). The x-axis indicates the proportion of different sites: p-distance. Each dot shows the amino acid sequence p-distance between the novel and known virus species. The International Committee on Taxonomy of Viruses species demarcation criteria are shown as orange dotted lines: greater than 0.3 in polyprotein, P1, and 2C+3CD regions for hepatoviruses (A), greater than 0.3 in polyprotein, P1 regions and 0.2 in 2C+3CD region for parechoviruses (B), and greater than 0.31 in the NS3 region and 0.31-0.36 in the NS5B region for pegiviruses (C).

**Supplemental Figure 3.**
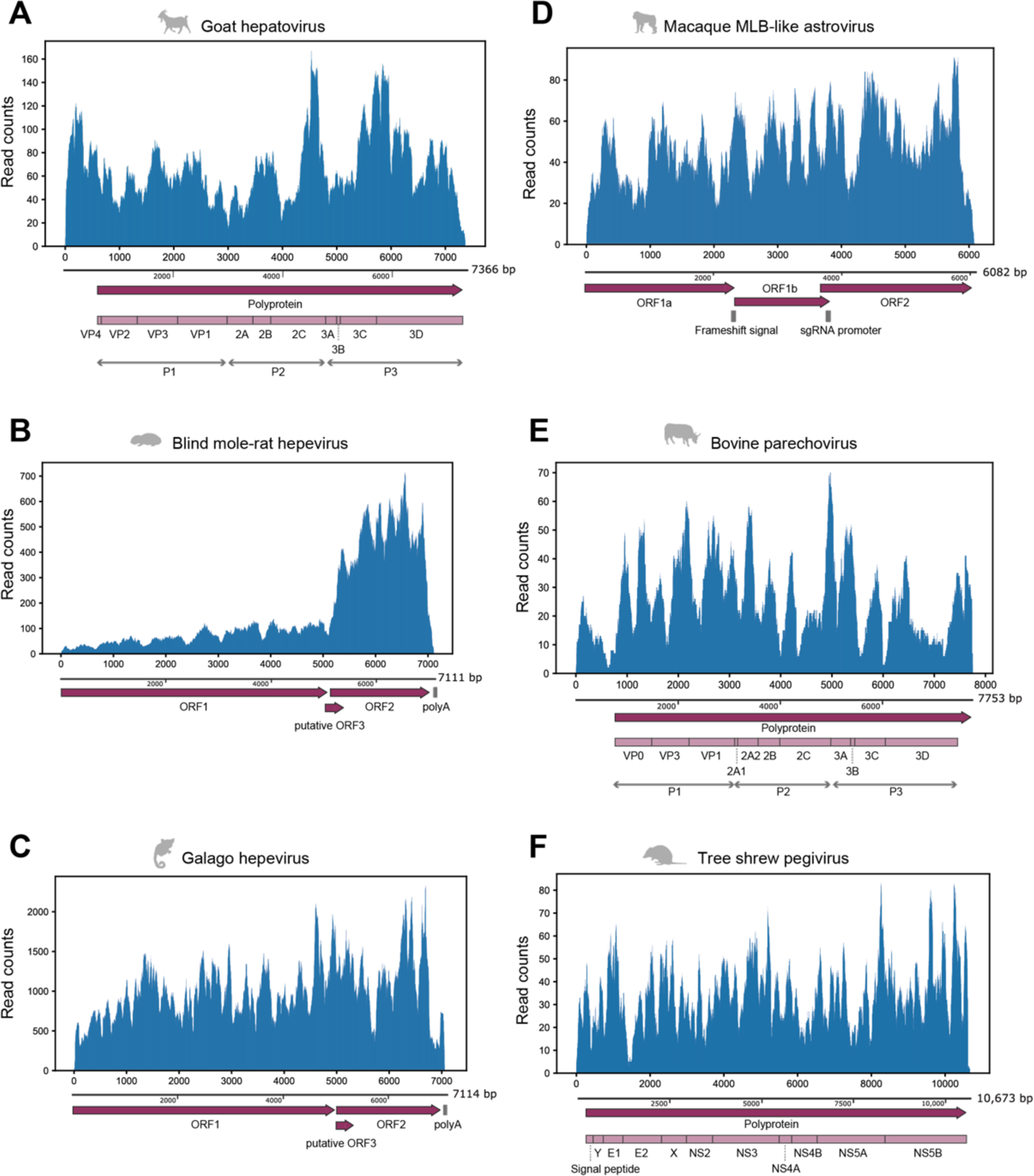
Mapping analysis using RNA-seq data in which the full-length viral genome was identified. (A-F) Read distributions mapped to the viral sequence: goat hepatovirus (A), blind mole-rat hepevirus (B), galago hepevirus (C), macaque MLB-like astrovirus (D), bovine parechovirus (E), and tree shrew pegivirus (F). The upper panel shows the virus genomic positions (x-axis) and read counts at each position (y-axis). The lower panel shows genomic annotations, such as protein-coding regions or signal sequences. Dark purple arrows indicate open reading frames (ORFs) in the viral genome. Light purple boxes show mature proteins predicted based on aligned positions with reference viruses (**details in Materials and Methods**). Gray vertical lines indicate nucleotide sequence features, such as polyadenylation signal (poly-A), ribosomal frameshift signal (frameshift signal), and promoter sequence for subgenomic RNA synthesis (sgRNA promoter).

## Supplemental Materials

**Supplemental Dataset 1. List of Sequence Read Archive run accession numbers, genome file, and sequence assembly method.**

**Supplemental Dataset 2. Information on RNA-seq data from experimental infection with viruses.**

**Supplemental Dataset 3. Information on possible viral contamination excluded from the totalization.**

**Supplemental Dataset 4. Information on manual curation for virus-host relationships.**

**Supplemental Dataset 5. Accession numbers of viral sequences used for phylogenetic analyses, viral genomic annotations, and comparing the International Committee on Taxonomy of Viruses species demarcation criteria.**

**Supplemental Dataset 6. Sequence Read Archive run accessions used for mapping analyses.**

**Supplemental Dataset 7. Sample metadata in which the goat hepatoviral infections were detected.**

**Supplemental Dataset 8. Sample metadata in which the blind mole-rat hepeviral infections were detected.**

**Supplemental Dataset 9. Sample metadata in which the galago hepeviral infections were detected.**

**Supplemental Dataset 10. Sample metadata in which the macaque MLB-like astrovirus infections were detected.**

**Supplemental Dataset 11. Sample metadata in which the bovine parechovirus infections were detected.**

**Supplemental Dataset 12. Sample metadata in which the tree shrew pegiviral infections were detected.**

**Supplemental Dataset 13. Bioinformatics tools and their versions used in this study.**

